# Mechanisms of trophic niche compression: evidence from landscape disturbance

**DOI:** 10.1101/329623

**Authors:** Francis J. Burdon, Angus R. McIntosh, Jon S. Harding

## Abstract

1. Natural and anthropogenic disturbances commonly alter patterns of biodiversity and ecosystem functioning. However, how food webs respond to these changes remains poorly understood. Here, we have described aquatic food webs using invertebrate and fish community composition, functional traits, and stable isotopes from twelve agricultural streams along a landscape disturbance gradient.
2. We predicted that excessive inputs of fine inorganic sediment (sedimentation) associated with agricultural land uses would negatively influence stream trophic diversity (e.g., reduced vertical and horizontal trophic niche breadths).
3. Food-web properties based on Bayesian analyses of stable isotope data (δ^13^C and δ^15^N) from consumers showed that increasing sediment disturbance was associated with reduced trophic diversity, indicated by the whole community (fish and invertebrates combined) occupying a smaller area in isotopic niche space. Reductions in trophic diversity were best explained by a narrowing of the consumer δ^13^C range, and to a lesser extent, consumer δ^15^N range along the sedimentation gradient.
4. We hypothesized that multiple mechanisms associated with sedimentation may have caused trophic niche ‘compression’. Decreased niche partitioning, driven by increasing habitat homogeneity, environmental filtering, and resource scarcity seemingly lead to a greater similarity in trophic roles. These pathways may have contributed to a reduction in trophic diversity, whereas increased resource homogeneity was seemingly less important.
5. Our results also indicate downward shifts in the vertical trophic position of benthic meospredators and invertebrate prey relative to higher consumers. This ‘trophic decoupling’ suggests that terrestrial resource subsidies may offset reductions of aquatic prey for larger stream fishes.
6. Sedimentation was associated with reduced trophic diversity, which may affect the functioning and stability of stream ecosystems. Our study helps explain how multiple mechanisms can influence food-web properties in response to this type of disturbance.

## Introduction

Food webs represent a holistic systems approach to characterizing patterns of biodiversity and energy flow by describing trophic interactions between consumers and resources (Thompson et al. 2012). One major goal in ecology is understanding how vertical (across-trophic level) and horizontal (within-trophic level) diversity contribute to food-web properties such as stability and energy transfer (Duffy et al. 2007, Thompson et al. 2012). Greater horizontal diversity may increase biomass and resource uptake (Duffy et al. 2007), and increased vertical diversity links to theory about food chain length and omnivory, with implications for trophic cascades (McHugh et al. 2010). To investigate trophic interactions in food webs, increasingly sophisticated methods are being used including stable isotopes, molecular tools, and network analyses (Raso et al. 2014, Wootton and Stouffer 2016). However, empirical studies of food webs that use underlying environmental gradients remain rare, despite their potential to help disentangle causal relationships between changing habitat conditions and ecosystem functioning (Thompson et al. 2012). We used a landscape disturbance gradient associated with agriculture-related inputs of fine sediment to streams to elucidate potential mechanisms causing changes in food-web structure and functioning.

Landscape disturbances associated with human activities are profoundly changing the natural dynamics of a wide range of systems (Turner 2010). In particular, the conversion of land for intensive agriculture has caused unprecedented landscape disturbance and habitat simplification, leading to species extinctions and the loss of ecosystem services (Tilman et al. 2001). One widespread and pervasive problem often associated with these activities is the degradation of stream habitat through inputs of fine inorganic sediment (Waters 1995). Excessive deposition of sediment (sedimentation), particularly in hard-bottomed streams, has been associated with species losses and threshold impacts on community composition (Matthaei et al. 2010, Burdon et al. 2013). With increasing sedimentation, reduced biomass and availability of basal resources and invertebrate prey may disrupt ‘bottom-up’ processes, thus affecting energy flow and species distributions in stream ecosystems (Osmundson et al. 2002, Suttle et al. 2004). Despite this, there have been few attempts to describe how sedimentation affects stream food-web properties, or fit this within a broader theoretical framework regarding trophic diversity and disturbance.

Stable isotope analysis (SIA) has been embraced as one of the main empirical tools for studying the trophic structure and dynamics of food webs (Layman et al. 2007a). Stable isotope ratios (typically carbon and nitrogen) are obtained from an organism’s tissues and represent cumulated energy from all trophic pathways in the food web leading to that individual. At the individual level, the trophic niche of an organism reflects its consumer-resource interactions, and thus represents dimensions of the Hutchinsonian ‘fundamental’ niche (i.e., the n-dimensional hypervolume) (Leibold 1995). The niche concept has been extended to a community-level property characterizing the trophic diversity and complexity of food webs using SIA (Layman et al. 2007a, Newsome et al. 2007). SIA has been widely used in food-web studies, ranging from the influences of habitat fragmentation on estuarine communities to the consequences of hydrodynamic disturbance in streams (Layman et al. 2007b, McHugh et al. 2010). Despite the promise of isotope-based metrics in describing the trophic structure and functioning of food webs (Layman et al. 2007a), it still remains poorly understood what underlying ecological processes contribute to the patterns these metrics help describe.

We predicted that there would be a ‘compression’ of the trophic niche (i.e., a reduction in trophic diversity as indicated by a smaller total area occupied by communities in isotopic niche space) in streams impacted by sedimentation. We were particularly interested in the horizontal and vertical niche breadths describing food-web structure. Sedimentation has wide-ranging impacts on stream ecosystems by homogenizing and degrading benthic habitat, thereby negatively affecting consumers and their resources (Waters 1995, Burdon et al. 2013). Thus, we predicted that the ‘realized niche’ breadths of stream food webs, as shown by the consumer δ^13^C and δ^15^N ranges (Layman et al. 2007a), would contract with increasing sedimentation. We hypothesized that four proximate mechanisms could explain changes in trophic diversity and realized niche breadths (Fig.1):

a. Niche ‘partitioning’ (Fig.1a), resulting from a heterogeneous resource base with specialists (Schoener 1974) that consume isotopically distinctive food sources, thus contributing to greater trophic diversity (i.e., niche differentiation). We expected that sites with low sediment would show the greatest taxonomic and trophic diversity owing to greater habitat and resource heterogeneity (Beisel et al. 2000);
b. Niche ‘elimination’ (Fig.1b), through habitat loss and degradation (i.e., environmental filtering; see Heino 2013) causing the non-random loss of specialist consumers. Here we expected to see reduced abundances and diversity of invertebrate ′predator′, ′shredder`, and ′grazer′ functional-feeding groups. For example, sedimentation has been associated with declines in grazing invertebrates reliant on hard substrates to consume diatoms (Rabení et al. 2005);
c. Niche ‘generalists’ (Fig.1c), where food scarcity leads to dietary convergence (i.e., increased polyphagy and omnivory), because all consumers become trophic generalists in accordance with optimal foraging theory (Pyke et al. 1977). We expected to see 1) little or no change in diversity, including consumer functional feeding groups, but changes in realized trophic niches with increasing sedimentation, and 2) an overall reduction in basal resources and invertebrate prey abundances consistent with sedimentation-induced resource scarcity; and
d. Niche ‘homogeneity’ (Fig.1d), where resource homogenization means consumers are more reliant on a narrower range of food items, because resources that are ‘hyper-abundant’ should be highly favored (Schoener 1974). Thus, we expected to see a numerical increase in invertebrates from the ′collector′ functional-feeding group correlated with an increase in fine benthic organic matter associated with sedimentation.

**Fig. 1.**
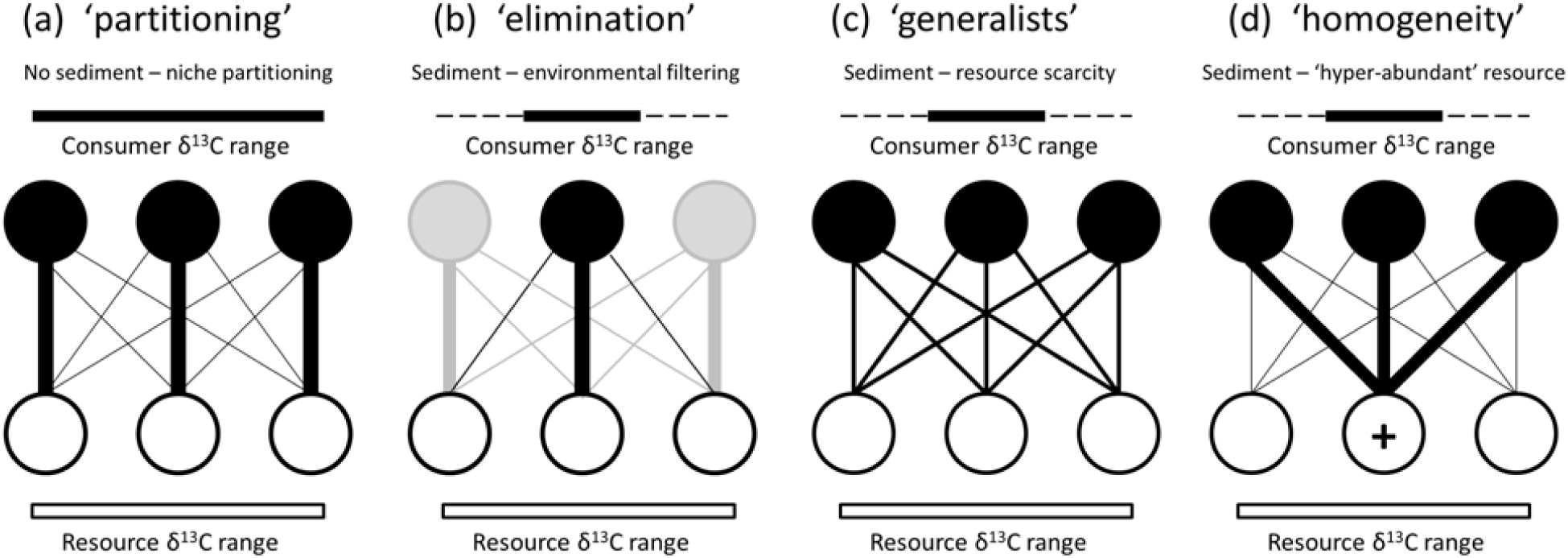
Four hypothetical consumer-resource interactions that may influence community trophic diversity (e.g., niche breadth). The mechanisms related to sediment disturbance are indicated. Although these examples describe changes in the consumer δ^13^C range (CR.B), the same principles apply to the consumer δ^15^N range (NR.B). It is assumed that interactions occur in ecosystems where basal resources show isotopic separation with a constant range, and consumer assimilation and fractionation rates are equivalent. (a) Consumer (solid circles) specialists concentrate interactions (black lines) on separate resources (open circles) thus ‘partitioning’ niche space and leading to an expansion of CR.B. (b) Niche ‘elimination’ where environmental filtering (i) removes specialist consumers (grey circles) as shown, or in an alternate scenario, (ii) removes or reduces preferred resources below functionally viable levels, causing interactions to cease. Both pathways lead to a contraction of CR.B. (c) Niche ‘generalists’, where resource scarcity leads to consumers equally sharing resources, causing greater trophic equivalence and a contraction of CR.B. (d) Resource ‘homogeneity’, where generalist and specialist consumers exploit an abundant resource (+ indicates an increase in resource availability), leading to a contraction of CR.B.

Niche processes (Fig. 1a-d) are nonexclusive, and all could occur simultaneously in real-world communities. We were interested in elucidating evidence for their relative contribution to changes in trophic diversity and realized niche breadths associated with sedimentation in stream ecosystems.

## Materials and Methods

### Study sites

We studied twelve 1^st^-3^rd^ order, wadeable, and perennially-flowing agricultural streams on the Canterbury Plains, located on the eastern side of New Zealand’s South Island. Streams were divided into six categories of deposited fine inorganic sediment (< 2 mm grain size; Burdon et al. 2013) and dissolved nutrients (i.e., a three by two factorial design incorporating differing levels of sediment and nutrients; Table 1). Independent stream reaches dominated by run-riffle sequences were 30-m long following Thompson and Townsend (2005). Reaches were sampled twice during base-flow conditions (habitat, resources, and invertebrates were sampled first, followed by fish communities) in the austral summer (6^th^ December 2009 to 5^th^ March 2010). For more information on study sites and site selection, see Appendix A.

**Table 1.**
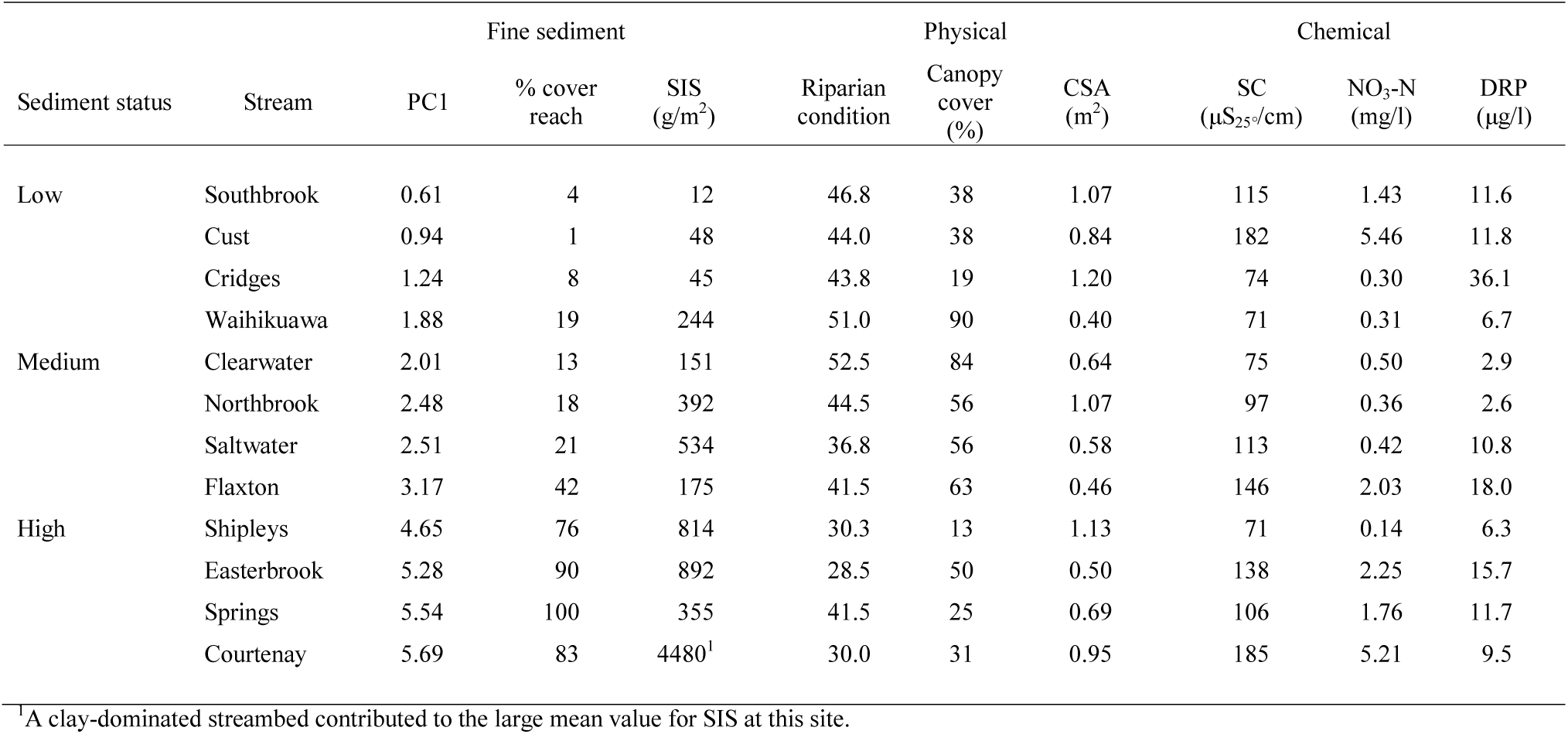
Description of 12 streams surveyed on the Canterbury Plains, South Island, New Zealand during the austral summer of 2009-2010. Streams are ranked along a sedimentation gradient (low to high). Additional site descriptors are provided in Appendix A, Table PC1, Principal Components Axis 1 scores of deposited sediment measures [+ constant (3)]; % cover reach, mean % sediment cover over the 30-m stream reach; SIS, mean suspendable inorganic sediment; CSA, stream cross-sectional area; SC, specific conductivity; DRP, dissolved reactive phosphorus.

### Data collection

To associate sedimentation-induced changes in benthic habitat with agricultural-related landscape disturbance, riparian habitat characteristics were recorded following Burdon et al. (2013). Briefly, riparian habitat quality was estimated for a 100-m stream segment (Richards et al. 1996) using a subjective index of 13 riparian attributes (Harding et al. 2009). Attributes were graded from poor (1) to excellent (5) on each bank, and bank scores were averaged and then summed to provide an index of total riparian condition (Table 1). Canopy cover was assessed using ocular estimation with a systematic approach (i.e., ten transects across the stream channel) to reduce variation and observer bias (Harding et al. 2009).

Deposited fine inorganic sediment (<1 mm grain size) was characterized using two subjective (visual) and two quantitative (physical) measurements following Burdon et al. (2013). To quantify sediment cover within each 30 m ‘reach’, ten 0.09 m^2^ gridded quadrats were stratified-randomly selected and substrate composition visually assessed at 25 grid intersections within each quadrat. Using the same methodology, substrate composition at the ‘patch’ scale was assessed in the five Surber samples collected for invertebrate communities (see below). Sediment depth was recorded at ten points in each reach using stratified-random selection. Suspendable inorganic sediment (SIS; g/m^2^) was similarly estimated from study reaches using the Quorer method (Clapcott et al. 2011, Burdon et al. 2013).

Water samples at each study site were collected on both sampling occasions for chemical analysis. Samples comprised of filtered (LabServ^®^ GF/F, 25 mm Ø; Thermo Fisher Scientific New Zealand Ltd, North Shore City, NZ) stream water (100 ml) collected in opaque acid-washed bottles, placed on ice, then frozen prior to analysis for nitrate (nitrate and nitrite nitrogen; mg/L) and DRP (dissolved reactive phosphorus; μg/L) using a SYSTEA Easychem discrete colorimetric auto-analyzer (SYSTEA S.p.A., Anagni, Italy; Appendix B.1.1). A further 1 L water sample was collected for estimation of total suspended solids (TSS; g/L) following Burdon et al. (2013).

Basal resources were collected from stream reaches for estimation of resource availability and stable isotope analyses. Using material collected in Surber samples, fine benthic and coarse particulate organic matter (FBOM and CPOM respectively; g/m^2^) was elutriated and separated using a 2-mm Endecott sieve. To estimate periphyton/biofilm availability (benthic AFDM mg/cm^2^ and chlorophyll-*a* μg/cm^2^), five cobbles or sediment samples were randomly collected from study reaches. Where fine sediment occurred in large deposits, a glass petri dish (45 mm Ø) was used to remove a circular section of sediment from the stream bed to 7.5 mm deep (Biggs and Kilroy 2000). Cobbles or sediment samples were immersed in 100% ethanol for 24 hours (in the dark) at 10° C for pigment extraction. Stone surface area and spectrophotometric chlorophyll-*a* was analyzed using methods in Biggs and Kilroy (2000). Filtered sub-samples (LabServ^®^ GF/C, 47 mm Ø) of periphyton/biofilm (brushed from stones in ethanol) and organic matter samples were dried (48 hours, 50° C), weighed, ashed (4 hours, 550° C), and re-weighed.

For stable isotopes analysis (SIA) of basal resources, we retained sub-samples of organic matter collected in the Surber samples (fine benthic organic matter, FBOM; and coarse particulate organic matter, CPOM). Algae and macrophyte samples were collected in the field, and biofilm samples were scraped from additional cobbles (*n* = 10) using a scalpel. Suspended fine particulate matter (SFPM) was collected in the channel thalweg using a plankton net (45 μm mesh) suspended in the water column for ten minutes. All samples were dried (48 hours, 50° C) prior to preparation for SIA.

To quantitatively sample invertebrate communities, five Surber samples (0.0625 m^2^, 250-μm mesh) were collected from channel thalwegs at stratified (evenly-spaced) randomly selected locations. Additional composite qualitative kicknet samples (250-μm mesh) were collected to encompass all microhabitats (e.g., riffles, macrophytes, marginal vegetation, wood, and leaf litter) present in the reach; invertebrate samples were placed on ice before being frozen. Kicknet samples were also used to obtain invertebrate specimens for SIA (see below). In the laboratory, thawed benthic samples were passed through a 500-µm Endecott sieve (Endecotts Limited, London, UK) and all invertebrates removed, identified, and counted to the lowest practicable level (usually genus) using invertebrate identification guides (e.g., Winterbourn et al. 2006). Taxa presence data from kick-nets were combined with Surber data to estimate total invertebrate taxonomic richness.

We additionally used functional traits (e.g., functional-feeding groups, FFGs) to help describe changes to invertebrate communities. FFGs are a well-established method to describe the functional roles of stream invertebrates (Cummins 1973). Based on invertebrate trait data from the New Zealand Freshwater Biodata Information System (FBIS), we assigned invertebrate trait affinity scores using ‘fuzzy-coding’ (Chevenet et al. 1994) to four FFGs: ‘predators’, ‘shredders’, ‘grazers’ and ‘collectors’. For more information about invertebrate trait analyses, see Appendix B.1.2, Table B1.

Fishes and decapod crustaceans were sampled using quantitative electro-fishing techniques (Appendix B.1.3) to estimate species richness and obtain specimens for stable isotopes analysis. At high sediment sites, sediment-induced turbidity reduced the effectiveness of our quantitative methods, thus only qualitative data has been used for fish community analyses.

### Stable isotope analysis

Although there are well-documented limitations of SIA, it remains one of the main tools to empirically describe food webs (Layman et al. 2007a). Stable isotope analysis (SIA) using carbon and nitrogen was performed on consumers and basal resources to assess changes to food web trophic structure along the sedimentation gradient. After thawing, composite kicknet samples were passed through a 500-μm sieve and retained stream invertebrates removed for isotopic analysis according to two criteria: highly abundant taxa (i.e., ≥ 100 individuals present in samples) and/or large bodied (i.e., > 5 mm body length); thus potentially contributing disproportionally to overall biomass. All fish taxa recorded from study streams were sampled for SIA, and where possible, three specimens of each species were collected to adequately characterize the range of sizes present at sites; dorsal muscle tissue was removed from thawed individuals for SIA.

To estimate isotopic content, samples of basal resources, invertebrates, and fish were dried (60 °C, 48 hours) and ground into a fine powder using a ceramic mortar and pestle. Stomach contents were removed from predatory invertebrates prior to drying (Jardine et al. 2005). Samples from individuals and aggregates of small-bodied invertebrate taxa (∼1.0 mg) and basal resources (∼2.0 mg) were then encapsulated into 8 × 5 mm tin capsules (OEA Laboratories Ltd., Cornwall, UK), and sent to the Stable Isotope Facility (University of California, Davis, CA, USA), where they were analyzed on a PDZ Europa 20–20 isotope ratio mass spectrometer (Sercon Ltd., Cheshire, UK). For more details about sample preparation and SIA methods, see Appendix B.1.4.

### Data analysis

Principal components analysis (PCA) was used to reduce our four measures of deposited fine sediment (‘reach’ and ‘patch’ cover, sediment depth, SIS) into a single sediment index explaining 85% of variation (PC1, Appendix B.2.1, Table B2). A constant (3) was added to PC1 to aid interpretability.

To evaluate trophic diversity, we used seven “community-wide” isotope-based metrics that describe different facets of trophic architecture (Table 2). The first four metrics measure the total extent of spacing within δ^13^C-δ^15^N bi-plot space, thus representing community-wide measures of trophic diversity; another two metrics reflect the relative position of species within niche space (Layman et al. 2007a). Another measure of trophic diversity was calculated using the standard ellipse area; an approach that is less susceptible to the effects of sample size and extreme values (Jackson et al. 2011). These metrics were calculated for ‘whole’ communities (fish and invertebrates combined), and individual communities (fish and invertebrates separately). Isotopic variability was also evaluated for basal resources (Appendix B.2.2).

**Table 2.**
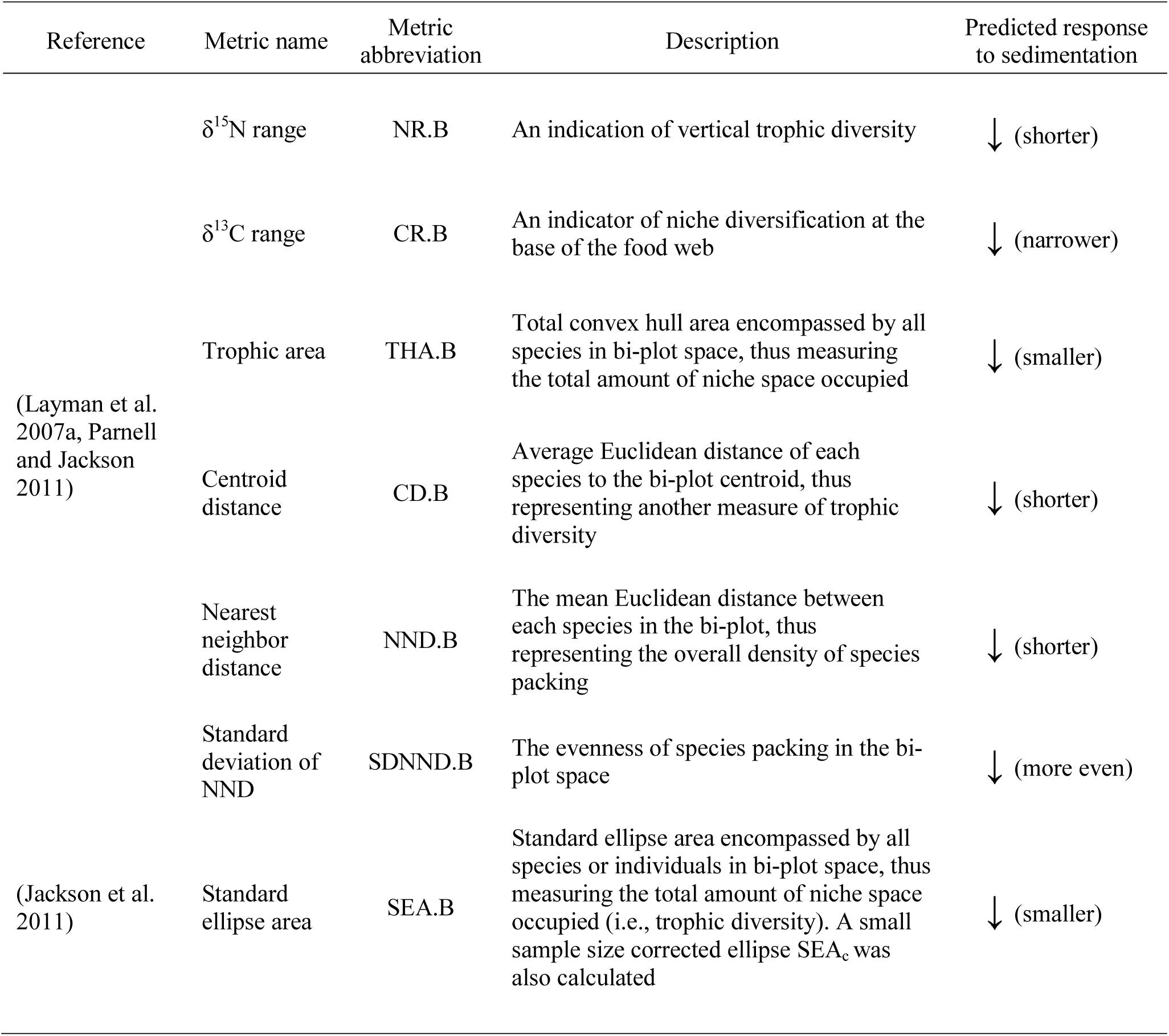
Stable isotope (δ^13^C and δ^15^N) metrics used to describe stream communities. The predicted responses of the metrics to sedimentation are also indicated.

We calculated the SIA community metrics from consumer isotope data using the “siar” package in R (Parnell and Jackson 2011). Following Bayesian inference techniques, the modes and relevant confidence intervals (i.e., distributions of estimated values) of the Layman metrics and standard ellipse areas were extracted from 100,000 posterior draws (Parnell and Jackson 2011). Community composition data were analyzed to better explain possible proximate changes influencing the SIA food-web metrics. Invertebrate taxa richness was rarefied to standardize richness estimates for samples of 1000 individuals using the ‘rarefy’ function in R. Fish community occupancy data (presence-absence) and relative abundances, invertebrate densities (individuals/0.0625 m^2^), and total and SIA invertebrate occupancy data were analyzed with non-metric multi-dimensional scaling (NMDS). Variation partitioning of community composition with sediment and nutrients was assessed using partial redundancy analyses (*p*RDA; Appendix B.2.4). All community analyses used the “vegan” package in R (Oksanen et al. 2013).

To assess the shape and significance of the relationship between deposited sediment and measured responses, regression models were selected using an information-theoretic approach (Burnham and Anderson 2002). Eight possible regression models were assessed. These were: linear, logarithmic, power, exponential, quadratic, asymptotic exponential, 3-parameter logistic, and 4-parameter sigmoidal curves. The lowest corrected Akaike Information Criterion (AIC_c_) values were used to identify the best-fitting model. Parameter estimates for the models shown in Figures 2-4 & 6 are reported in Table 3, and in Appendix C for all other regressions.

**Table 3.**
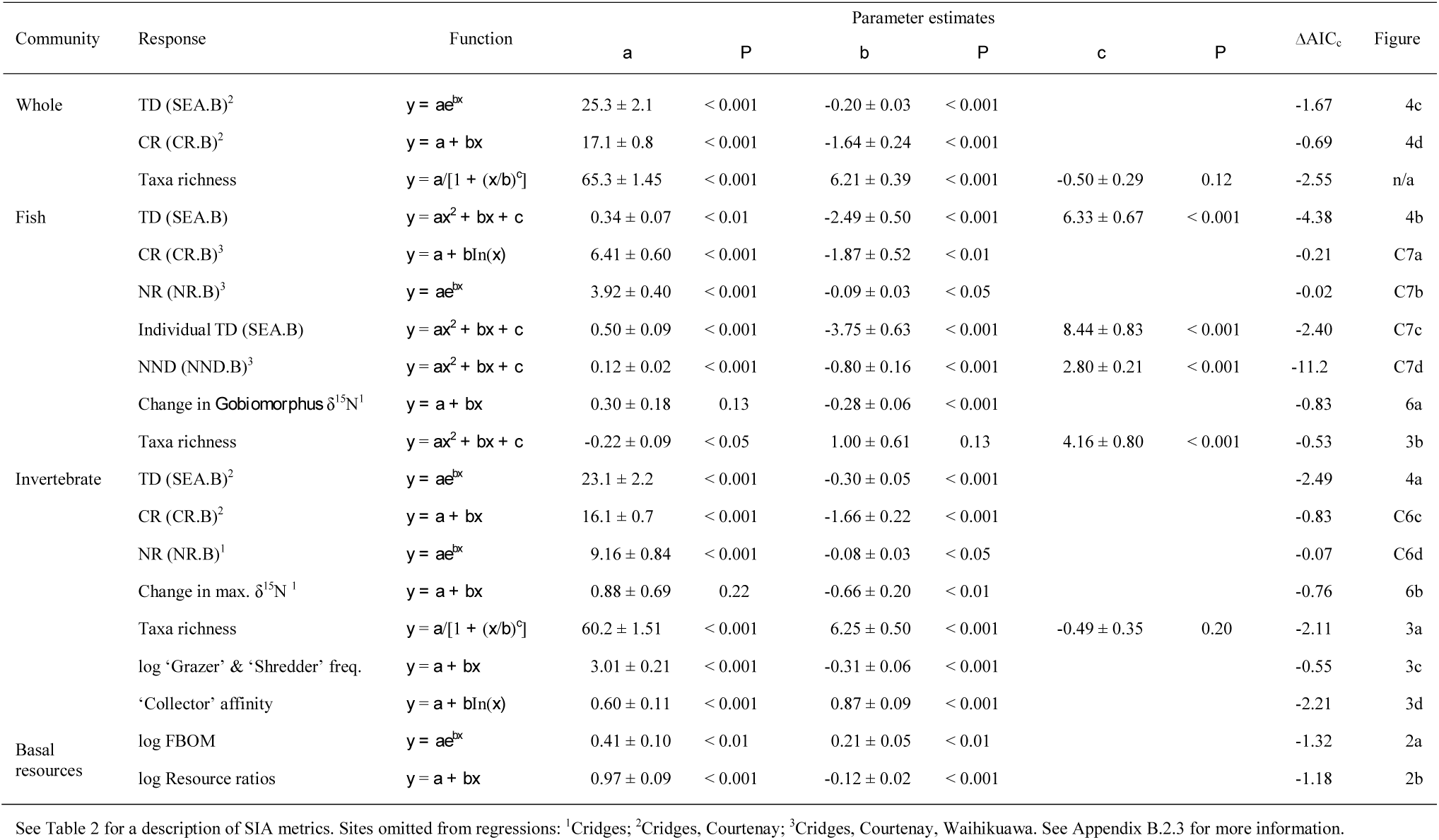
Parameter estimates for significant regressions of sediment (PC1) against response variables including food-web properties (whole, fish and invertebrate community metrics using stable isotopes of δ^13^C and δ^15^N), taxa richness, invertebrate functional-feeding groups, and basal resources. FBOM, fine benthic organic matter. ΔAIC_c_ (i.e., AIC_c_^i^ – AIC_c_^j^, where *i* is the minimum AIC_c_ recorded and *j* is the next lowest model AIC_c_ value).

| Community | Response | Function | Parameter estimates | | | | | | $\Delta\text{AIC}_c$ | Figure |
| --- | --- | --- | --- | --- | --- | --- | --- | --- | --- | --- |
| | | | $a$ | $P$ | $b$ | $P$ | $c$ | $P$ | | |
| Whole | TD (SEA.B)<sup>2</sup> | $y = ae^{bx}$ | $25.3 \pm 2.1$ | $< 0.001$ | $-0.20 \pm 0.03$ | $< 0.001$ | | | -1.67 | 4c |
| | CR (CR.B)<sup>2</sup> | $y = a + bx$ | $17.1 \pm 0.8$ | $< 0.001$ | $-1.64 \pm 0.24$ | $< 0.001$ | | | -0.69 | 4d |
| | Taxa richness | $y = a/[1 + (x/b)^c]$ | $65.3 \pm 1.45$ | $< 0.001$ | $6.21 \pm 0.39$ | $< 0.001$ | $-0.50 \pm 0.29$ | 0.12 | -2.55 | n/a |
| Fish | TD (SEA.B) | $y = ax^2 + bx + c$ | $0.34 \pm 0.07$ | $< 0.01$ | $-2.49 \pm 0.50$ | $< 0.001$ | $6.33 \pm 0.67$ | $< 0.001$ | -4.38 | 4b |
| | CR (CR.B)<sup>3</sup> | $y = a + b\ln(x)$ | $6.41 \pm 0.60$ | $< 0.001$ | $-1.87 \pm 0.52$ | $< 0.01$ | | | -0.21 | C7a |
| | NR (NR.B)<sup>3</sup> | $y = ae^{bx}$ | $3.92 \pm 0.40$ | $< 0.001$ | $-0.09 \pm 0.03$ | $< 0.05$ | | | -0.02 | C7b |
| | Individual TD (SEA.B) | $y = ax^2 + bx + c$ | $0.50 \pm 0.09$ | $< 0.001$ | $-3.75 \pm 0.63$ | $< 0.001$ | $8.44 \pm 0.83$ | $< 0.001$ | -2.40 | C7c |
| | NND (NND.B)<sup>3</sup> | $y = ax^2 + bx + c$ | $0.12 \pm 0.02$ | $< 0.001$ | $-0.80 \pm 0.16$ | $< 0.001$ | $2.80 \pm 0.21$ | $< 0.001$ | -11.2 | C7d |
| | Change in *Gobiomorphus* $\delta^{15}\text{N}$<sup>1</sup> | $y = a + bx$ | $0.30 \pm 0.18$ | 0.13 | $-0.28 \pm 0.06$ | $< 0.001$ | | | -0.83 | 6a |
| | Taxa richness | $y = ax^2 + bx + c$ | $-0.22 \pm 0.09$ | $< 0.05$ | $1.00 \pm 0.61$ | 0.13 | $4.16 \pm 0.80$ | $< 0.001$ | -0.53 | 3b |
| Invertebrate | TD (SEA.B)<sup>2</sup> | $y = ae^{bx}$ | $23.1 \pm 2.2$ | $< 0.001$ | $-0.30 \pm 0.05$ | $< 0.001$ | | | -2.49 | 4a |
| | CR (CR.B)<sup>2</sup> | $y = a + bx$ | $16.1 \pm 0.7$ | $< 0.001$ | $-1.66 \pm 0.22$ | $< 0.001$ | | | -0.83 | C6c |
| | NR (NR.B)<sup>1</sup> | $y = ae^{bx}$ | $9.16 \pm 0.84$ | $< 0.001$ | $-0.08 \pm 0.03$ | $< 0.05$ | | | -0.07 | C6d |
| | Change in max. $\delta^{15}\text{N}$<sup>1</sup> | $y = a + bx$ | $0.88 \pm 0.69$ | 0.22 | $-0.66 \pm 0.20$ | $< 0.01$ | | | -0.76 | 6b |
| | Taxa richness | $y = a/[1 + (x/b)^c]$ | $60.2 \pm 1.51$ | $< 0.001$ | $6.25 \pm 0.50$ | $< 0.001$ | $-0.49 \pm 0.35$ | 0.20 | -2.11 | 3a |
| | log 'Grazer' & 'Shredder' freq. | $y = a + bx$ | $3.01 \pm 0.21$ | $< 0.001$ | $-0.31 \pm 0.06$ | $< 0.001$ | | | -0.55 | 3c |
| | 'Collector' affinity | $y = a + b\ln(x)$ | $0.60 \pm 0.11$ | $< 0.001$ | $0.87 \pm 0.09$ | $< 0.001$ | | | -2.21 | 3d |
| Basal resources | log FBOM | $y = ae^{bx}$ | $0.41 \pm 0.10$ | $< 0.01$ | $0.21 \pm 0.05$ | $< 0.01$ | | | -1.32 | 2a |
| | log Resource ratios | $y = a + bx$ | $0.97 \pm 0.09$ | $< 0.001$ | $-0.12 \pm 0.02$ | $< 0.001$ | | | -1.18 | 2b |

**Fig. 2.**
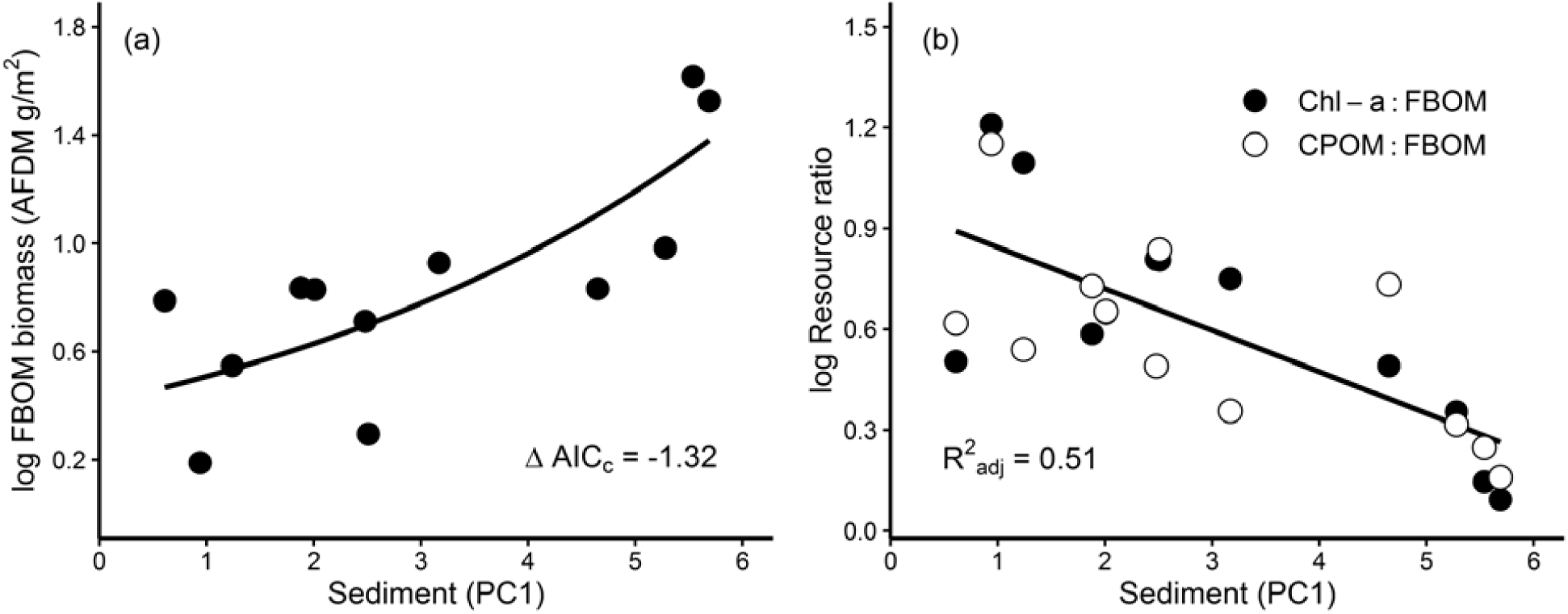
Relationship between sediment (PC1) and: (a) log_10_ (*x* + 1) fine benthic organic matter (FBOM; ash-free dry mass g/m^2^), and (b) log_10_ (*x* + 1) ratio of chlorophyll-a concentrations (μg/cm^2^) and coarse-particulate organic matter (CPOM; ash-free dry mass, g/m^2^) to FBOM. For parameter estimates, see Table 3.

Two sites deviated from the general patterns documented by our whole community SIA analyses and were treated as outliers. One site (Courtenay) seemingly had a novel carbon source (e.g., methanogenesis) not utilized at other sites, and the other site (Cridges) had an apparent nutrient disturbance prior to sampling, leading to a disequilibrium in consumer isotopic signatures. An additional site (Waihikuawa) was excluded from the regression analyses of the fish community involving the ‘Layman’ metrics due to the low taxa richness (< 3). For more information on the selection of regression models and the treatment of outliers, see Appendix B.2.3. We conducted all analyses in the R language and environment for statistical computing (R Core Team 2015).

## Results

### Landscape disturbance gradient

Stream sedimentation was more likely to occur with upstream riparian habitat degradation, as evidenced by the correlation of our deposited sediment index (PC1) with decreasing riparian condition (Table 1; Pearson’s product-moment correlation, *r* = −0.749, *t_10_* = −3.58, *P* < 0.01). Canopy cover decreased with riparian degradation (*r* = 0.568, *t_10_* = 2.18, *P* = 0.054) and increasing stream size (*r* = −0.713, *t_10_* = 3.22, *P* < 0.01), but not with sediment (*r* = −0.308, *t_10_* = - 1.03, *P* = 0.330). Other factors, such as stream size and nutrient concentrations, were not associated with riparian condition or sediment (Table 1).

### Basal resources

SIA was performed on basal resources to determine if variation in their isotopic signatures potentially contributed to the patterns in consumer trophic diversity described below. The number of basal resource types used for SIA did not change with increasing sedimentation (*r* = 0.433, *t_8_* = 1.57, *P* = 0.155). Likewise, there was no change in the basal resource δ^13^C range (*r* = 0.224, *t_8_* = 0.650, *P* = 0.534) and coefficient of variation (*r* = −0.281, *t_8_* = −0.828, *P* = 0.431) along the sediment gradient, meaning observed changes in consumer isotopic niches were unlikely caused by differences in the isotopic signatures of basal resources. The same patterns were observed when including macrophytes; for the full description of results relating to the basal resource SIA metrics, see Appendix C.1, Tables C1-2.

However, increasing deposited sediment (PC1) was associated with changes in the standing stocks of basal resources. The biomass of fine benthic organic matter (FBOM) increased with sediment (Table 3, Fig. 2a), whereas primary producers (benthic chlorophyll-*a* concentrations; *r* = −0.439, *t_10_* = −1.547, *P* = 0.153) and terrestrial detritus (coarse particulate organic matter, CPOM, AFDM g/m^2^; *r* = 0.139, *t_10_* = 0.445, *P* = 0.666) did not change significantly. This meant the ratio of benthic chlorophyll-*a* and CPOM to FBOM decreased with increasing sedimentation (Homogeneity of lines test: *t_1,20_* = 0.80, *P* = 0.44; *F_1,22_* = 25.1, *P* < 0.001, *R^2^* = 0.512, Fig. 2b). This scenario was more consistent with the conditions required for niche ′homogeneity` (i.e., resource homogenization) than niche ′generalists′ (i.e., resource scarcity).

### Invertebrate community responses

Increasing deposited sediment (PC1) was associated with reductions in invertebrate community taxonomic richness (Table 3, Fig. 3a). However, rarefied invertebrate taxa richness did not change significantly with sediment (Logarithmic regression, ?AICc = −0.16, *F_1,10_* = 5.68, *P* = 0.173, *R^2^* = 0.178), suggesting that lower invertebrate abundances contributed to reduced richness. Sediment alone significantly explained variation in invertebrate community structure (e.g., 27% of variation using abundances, *p*RDA, *F_1,9_* = 4.57, *P* < 0.01), whereas the joint and independent contributions of nutrients (phosphorus and nitrate) had no significant influence (Appendix C.2, Table C5, Fig. C2).

**Fig. 3.**
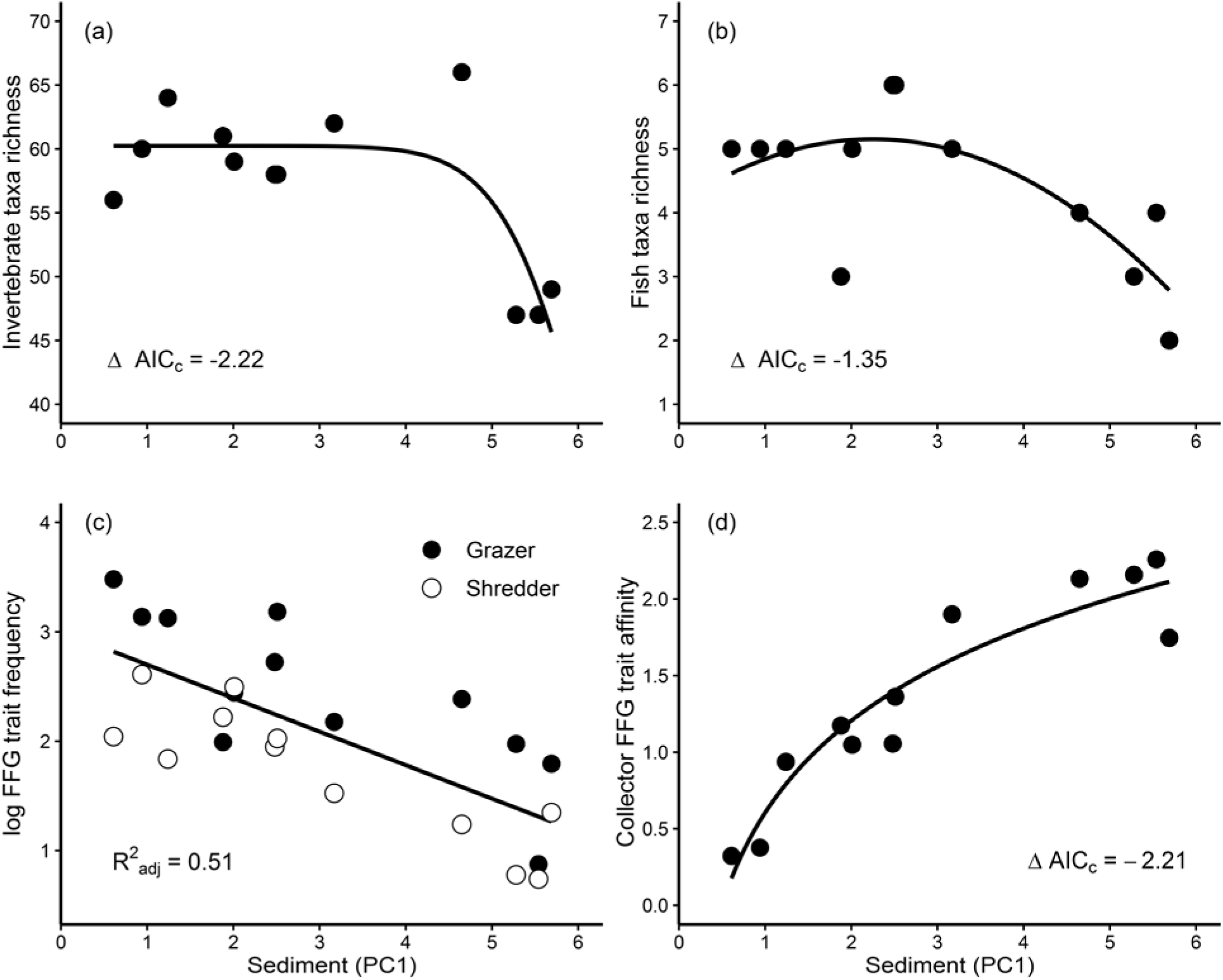
Structural and functional biodiversity changes: relationships between deposited sediment (PC1) and: (a) invertebrate taxa richness, (b) fish taxa richness, (c) log_10_ invertebrate ‘grazer’ and ‘shredder’ functional-feeding group (FFG) trait frequencies, and (d) ‘collector’ FFG trait affinity (FFG trait frequency per individual). For parameter estimates, see Table 3.

Deposited sediment (PC1) was associated with changes in invertebrate functional-feeding groups (FFGs) that were consistent with the niche ‘elimination’ mechanism. Using community abundance data, the frequency of ‘grazers’ and ‘shredders’ significantly decreased with sediment (Homogeneity of lines test: *t_1,20_* = 0.34, *P* = 0.74; *F_1,22_* = 25.0, *P* < 0.001, *R^2^* = 0.511, Fig. 3c). Significant negative associations were also seen using total occupancy (i.e., presence) data for specialist ‘grazers’ (*r* = −0.616, *t_10_* = −2.47, *P* < 0.05) and ‘shredders’ (*r* = −0.601, *t_10_* = −2.38, *P* < 0.05). There was evidence using invertebrate FFGs that refuted the ′niche homogeneity` mechanism. Despite the increase in FBOM, the frequency of ‘collectors’ did not significantly change with sediment (*r* = 0.255, *t_10_* = 0.835, *P* = 0.423). However, the trait affinity (mean trait score per individual) for ′collectors′ increased asymptotically (Table 3, Fig. 3d) with sediment, further supporting the niche ‘generalists’ mechanism. This indicated that more generalist consumers (‘collectors’) formed the core of invertebrate food webs as more specialist ‘grazers’ and ‘shredders’ were lost with sedimentation. For a summary of all invertebrate functional trait results, see Appendix C, Table C4.

These changes in invertebrate FFGs could be associated with the influence of sediment on invertebrate community SIA metrics. The trophic diversity (TD) of invertebrate communities, as measured by the Bayesian standard ellipse area (SEA.B) declined with increasing levels of deposited sediment (Table 3, Fig. 4a), reflecting changes in both horizontal and vertical trophic diversity. Sedimentation had a negative influence on the invertebrate δ^13^C carbon range (CR.B; *F_1,8_* = 55.0, *P* < 0.001,*R*^2^_adj_; Appendix C, Fig. C6c) and δ^15^N nitrogen range (NR.B, Table 3; Appendix C, Fig. C6d). Changes in invertebrate community SIA metrics were not confounded by reduced taxonomic richness, because there was no significant difference in the numbers of taxa used for SIA along the sediment (PC1) gradient (Pearson’s product-moment correlation, *r* = −0.231, *t_10_* = −0.751, *P* = 0.470). Isotopic metrics and abbreviations are further explained in Table 2.

**Fig. 4.**
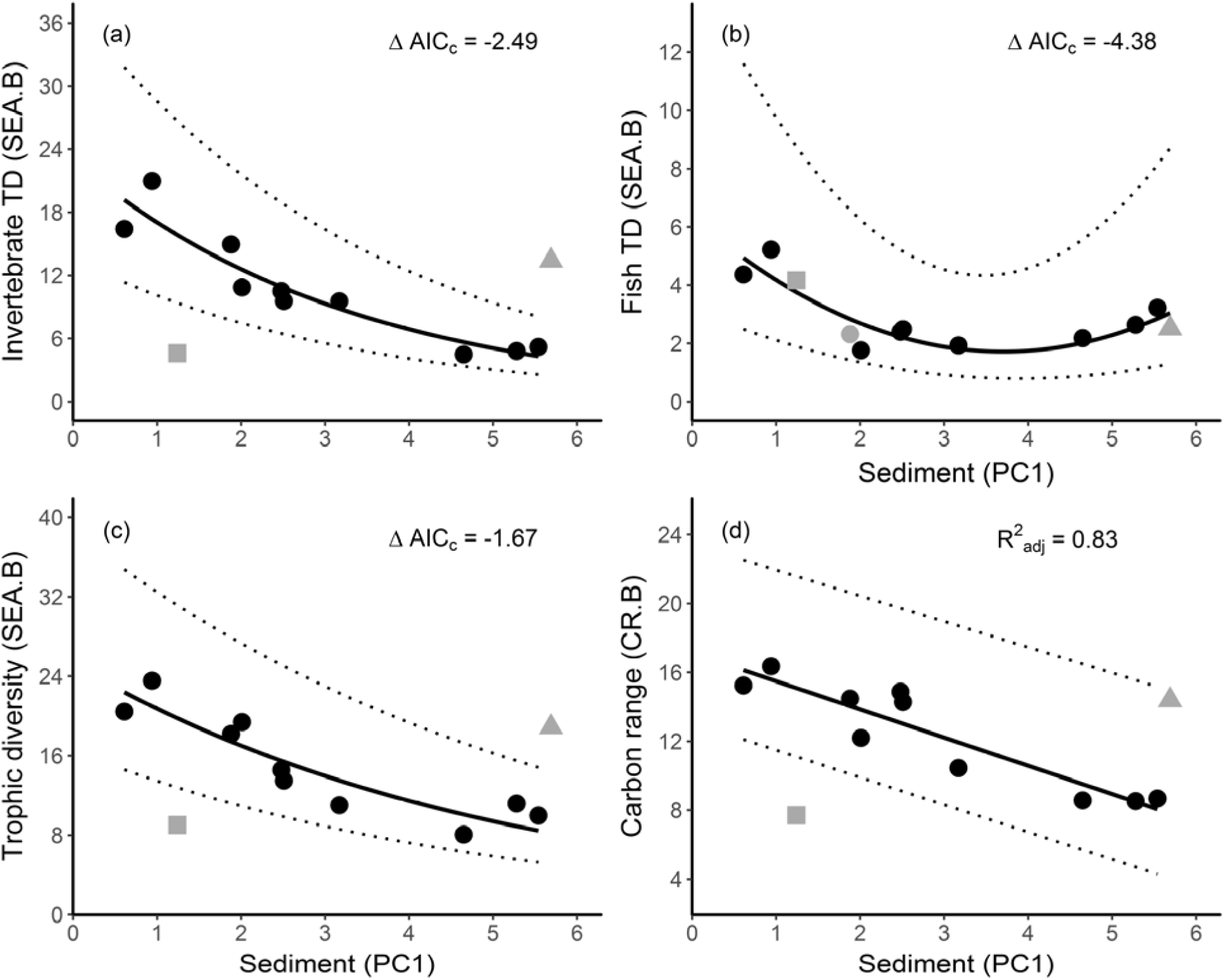
Trophic niche ‘compression’ and ‘expansion’: Results from Bayesian community stable isotope (δ^13^C and δ^15^N) metrics of twelve agricultural streams across a sedimentation gradient. Relationship between deposited sediment and: (a) whole community (fish and invertebrates) trophic diversity (SEA.B), (b) whole community consumer δ^13^C range (CR.B), (c) invertebrate trophic diversity (SEA.B), and (d) fish trophic diversity (SEA.B). For parameter estimates, see Table 3. Dotted lines indicate distributions of estimated values (i.e., confidence intervals) of 95% (a,c), 80% (c), and 90% (d). Outlier sites shown in grey: square (Cridges), triangle (Courtenay), circle (Waihikuawa); see Appendix B.2.3 for more information.

### Fish community responses

Fish community richness showed a weak unimodal relationship with sediment, but richness was generally lower at high levels of sediment compared to low or intermediate levels (Table 3, Fig. 3b). However, redundancy analysis indicated that neither sediment nor nutrients significantly explained fish community structure using occupancy data (Appendix C, Table C5).

In contrast, fish communities, as measured by SIA metrics, were influenced by sedimentation (PC1). Fish community TD (SEA.B) had a ‘U-shaped’ relationship with increasing sediment using species means (Table 3, Fig. 4b); an identical pattern was observed using individual fish isotope data (i.e., not species means; Table 3; Appendix C, Fig. C7c). This was likely influenced by reduced species packing (NND.B) at low and high sediment sites, leading to a similarly ‘U-shaped’ relationship with increasing sediment (Table 3; Appendix C, Fig. C7d). Fish community δ^15^N ranges (NR.B, Table 3) and δ^13^C ranges (CR.B; Logistic regression, *F_1,7_* = 9.47, *P* < 0.05,*R*^2^_adj_) both declined, albeit weakly, in a non-linear fashion with sedimentation (Appendix C, Fig. C7a,b). The general ‘decay’ pattern of these relationships indicated that changes in the isotopic ranges for fish communities were nearly invariant over much of the sediment gradient (i.e., most of the change occurred between low and intermediate sediment levels).

### Whole community food-web responses

The trophic diversity (TD) of the whole community (invertebrates and fish) was reduced by sedimentation (e.g., Fig. 5). Whole community TD (SEA.B) was negatively associated with deposited sediment (Table 3, Fig. 4c). This reduction was strongly linked to changes in horizontal, but not vertical diversity. Whole communities showed narrowing consumer δ^13^C carbon ranges (CR.B) with increasing sediment (*F_1,8_* = 45.8, *P* < 0.001, *R*^2^_adj_); Table 3, Fig. 4d), whereas the nitrogen range (NR.B; *r* = 0.076, *t_8_* = 0.216, *P* = 0.835) did not change along the sediment gradient (Appendix C, Table C6, Fig. C6b).

**Fig. 5.**
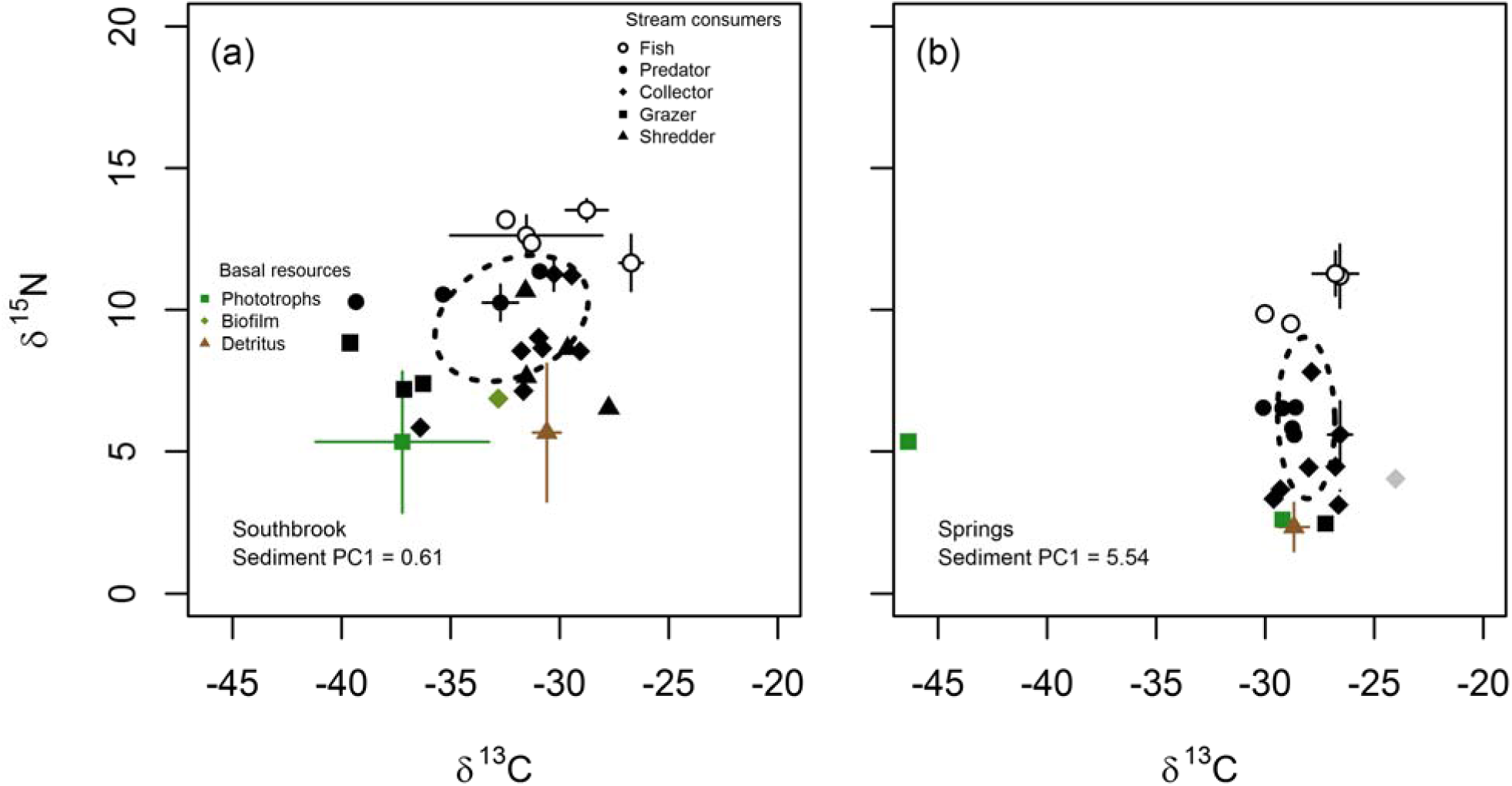
Stable isotope biplots of δ^13^C ‰ (^13^C:^12^C) and δ^15^N ‰ (^15^N:^14^N) describing food webs from two streams influenced by (a) low and (b) high levels of deposited sediment (PC1). Symbols in solid black indicate invertebrate consumers and their respective functional-feeding groups. Black-dashed line, standard ellipse SEA_c_ (see Table 2). Where appropriate, errors bars (± 1 standard deviation) for consumers and basal resources are shown. The grey diamond in (b) indicates an invertebrate consumer (the ostracod *Herpetocypris pascheri*) that was excluded from analyses due to ^13^C-enriched values (a potential artefact of exoskeleton chemistry). The green square near the y-axis in plot (b) represents a bryophyte; the extremely depleted δ^13^C value may have been influenced by methanogenesis. See Appendix C, Figs. C3-5 for stable isotope biplots from all sites.

Interestingly, the stable isotope signatures of benthic mesopredators (i.e., middle trophic-level predators which both predate and are predated upon) were influenced by sedimentation relative to their invertebrate prey and potential piscivorous predators. The amount of change in mean δ^15^N for the benthic fish *Gobiomorphus cotidianus* and *G. breviceps* relative to the mean δ^15^N for larger stream fish (i.e., salmonids and aguilliformes) increased negatively along the sediment gradient (Homogeneity of lines test: *t_1,14_* = 0.73, *P* = 0.48; *F_1,14_* = 23.3, *P* < 0.001, *R^2^* = 0.598, Fig. 6a). There was no relationship between sediment and the larger stream fish δ^15^N (*r* = 0.134, *t_9_* = 0.407, *P* = 0.694). Likewise, the amount of change in the maximum δ^15^N values for stream invertebrates (i.e., the top predatory invertebrate) relative to the mean δ^15^N values for the *Gobiomorphus* spp. also increased negatively with increasing sedimentation (Homogeneity of lines test: *t_1,14_* = −0.13, *P* = 0.90; *F_1,15_* = 10.4, *P* < 0.01, *R^2^* = 0.369, Fig. 6b). This indicated that these benthic mesopredators tracked ′compressed′ invertebrate prey down, whereas larger stream fishes were able to maintain their trophic position, helping to explain the absence of change in vertical trophic diversity for the whole communities.

**Fig. 6.**
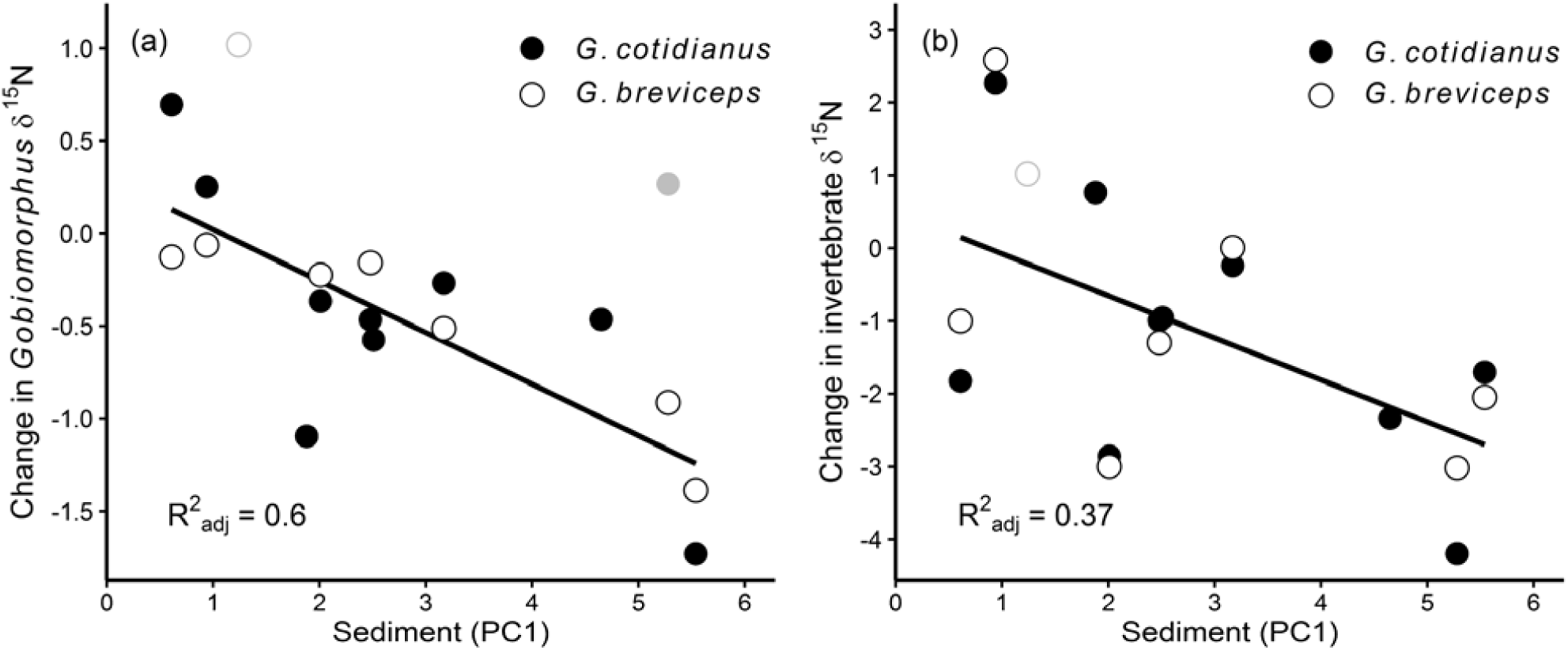
‘Trophic decoupling’ through niche compression: the relationship between deposited sediment and (a) change in the mean δ^15^N values for the stream fishes *Gobiomorphus* spp. relative to the mean δ^15^N values for larger stream fish (i.e., salmonids and anguilliformes); and (b) the change in the maximum δ^15^N values for stream invertebrates (i.e., the top predatory invertebrate) relative to the mean δ^15^N values for *Gobiomorphus* spp. For parameter estimates, see Table 3. Outlier sites shown in grey: *G. breviceps* (Cridges, open circle); in (a) *G. cotidianus* (Easterbrook, closed circle); for more information, see Appendix B.2.3

## Discussion

Our results reveal evidence of reduced trophic diversity in stream food webs along a landscape disturbance gradient, where sedimentation associated with degraded riparian habitat influenced the ‘compression’ of isotopic niche space. We postulated that four types of consumer-resource mechanisms may have contributed to the extent of niche breadth changes measured by the consumer δ^13^C and δ^15^N ranges (Fig. 1a-d). Greater niche partitioning at low sediment levels leading to wider consumer δ^13^C and δ^15^N ranges (Fig. 1a) appeared to be replaced by two mechanisms (i.e., niche ‘elimination’ and ‘generalists’) driving narrower consumer ranges in isotopic niche space (Fig. 1b,c). We further discuss the underlying ecological mechanisms driving trophic niche ‘compression’, and explore the consequences of this for the stability and functioning of food webs.

### Niche ‘partitioning’ and ‘elimination’ mechanisms

The niche ‘partitioning’ mechanism (Figs. 1a, 7a) reflects theories about niche differentiation, and is intimately linked to the relationship between habitat and species diversity (Cardinale 2011). Substrate heterogeneity in streams, for example, frequently has a strong positive correlation with invertebrate diversity, and this relationship is often attributed to the “greater number of niches” in areas of more heterogeneous habitat (Beisel et al. 2000). In contrast, sedimentation normally results in more homogenous instream habitat dominated by fine particles (Waters 1995, Burdon et al. 2013). This may have functional consequences, because larger substrate particles typically exhibit greater stability during high flows, accumulate more basal resources, and provide spatial refuge, thus generally resulting in greater abundances and diversity of biota (Quinn and Hickey 1990). These factors may explain why we saw greater trophic diversity in more heterogeneous sites with low levels of sedimentation.

Perturbations such as sedimentation may drive non-random (i.e., trait-mediated) losses in biodiversity that have functional consequences for food webs (Woodward 2009). In our example, sedimentation associated with landscape disturbance may have led to environmental filtering of sensitive taxa (akin to ‘species sorting’; Heino 2013), thus supporting the niche ‘elimination’ hypothesis (Figs. 1b, 7b). Sedimentation can exert stresses on stream invertebrates through dislodgement by ‘saltating’ fine sediment particles and habitat degradation (Waters 1995, Burdon et al. 2013). We saw substantial reductions in taxa richness with decreased frequencies of more specialized invertebrate consumers (i.e., ‘grazers’ and ‘shredders’) along the sedimentation gradient; results similar to other published studies (Rabení et al. 2005, Yule et al. 2010). We also associated sediment disturbance with the disappearance of specialist predators such as the rheophilic, free-living caddisfly *Hydrobiosis frater* (Trichoptera: Hydrobiosidae) based on occupancy data (*Author′s unpublished data*). These losses of specialist consumers likely contributed to the reduction of trophic diversity with increasing sedimentation. Similar to our study, isotopic niche contraction due to species extinctions has been observed in fragmented estuarine communities (Layman et al. 2007b), streams impacted by acid-mine drainage (Hogsden and Harding 2014), and primate communities facing multiple anthropogenic pressures (Crowley et al. 2012).

### Niche ‘generalists’ and ‘homogeneity’ mechanisms

Alternatively, species turnover along environmental gradients may help to regulate trophic structure in certain instances (Leibold et al. 1997). However, the ecological roles of generalist species able to persist in disturbed habitats can also be altered (Layman et al. 2007b, Lindo et al. 2012). Many stream consumers are polyphagous (Cummins 1973), and Yule et al. (2010) found that several invertebrate taxa which were ‘grazers’ at sites unaffected by sediment became facultative ‘collectors’ at impacted sites. Thus, increasing sedimentation may cause changes in the availability of resources, thus inducing shifts in consumer diets.

Optimal foraging theory predicts that food scarcity should lead to dietary convergence, because all species become generalists (Pyke et al. 1977). This mechanism underpins the most likely scenario for the niche ‘generalists’ hypothesis (Figs. 1c, 7b). In our study, resource abundances associated with grazing (chlorophyll-a) and shredding (CPOM) did not change along the sedimentation gradient, despite strong changes in invertebrate FFG associated with these resources. However, using standing stocks of biomass may be misleading because studies have indicated impaired transfers of energy in streams impacted by sediment through decreased resource quality and ‘shredding’ activity (Waters 1995, Burdon 2013). We saw indirect evidence of this with more ^13^C-enriched values of biofilms and CPOM along the sediment gradient (Appendix C, Fig. C1), indicative of greater heterotrophy consistent with burial by sediment. This change in consumer access to resources (i.e., scarcity) includes reduced prey abundances for higher predators, and may have contributed to the narrowing δ^13^C and δ^15^N ranges along the sediment gradient.

We observed an overall shortening of δ^15^N ranges for invertebrate and fish communities, but not the whole community. Part of this discrepancy was explained by ‘trophic decoupling’ (Figs. 6, 7b), where top predators and prey were ‘pulling apart’ in isotopic niche space along the sediment gradient. Here, top predators were able to maintain trophic position, potentially by feeding on terrestrial resource subsidies (see below), whereas benthic mesopredators and invertebrate prey were seemingly influenced by ‘niche compression’ mechanisms, thus explaining the absence of vertical diversity changes for the whole food web. Importantly, the shape of the response by the fish community δ^15^N ranges with increasing sediment was only a shallow decay, with the δ^15^N ranges invariant along the majority of the gradient. This response may have contributed to the increases in fish community TD at high sediment levels using the standard ellipse approach, despite narrowing δ^13^C and δ^15^N ranges. Standard ellipses are less susceptible to small sample sizes and extreme values (Jackson et al. 2011). In contrast, we did observe strong narrowing of δ^15^N ranges for invertebrate consumers along the sediment gradient. This was despite no obvious changes in the abundances of predatory invertebrates (*sensu* niche ‘elimination’), suggesting that predators may have become more omnivorous (e.g., through species-sorting, turnover, and dietary plasticity) in response to sediment-induced resource scarcity, thus acquiring δ^15^N signatures more similar to their prey. Consistent with these interpretations, Yule et al. (2010) found that increasing sedimentation in a tropical river generally led to simpler food webs with increased connectance, suggesting more generalized trophic roles. Other disturbances may cause resource scarcity and increasing trophic equivalence. McHugh et al. (2010) found evidence of realized omnivory varying systematically with hydrodynamic disturbance when considering the trophic position of stream fishes and abundances of predatory invertebrates.

**Fig. 7.**
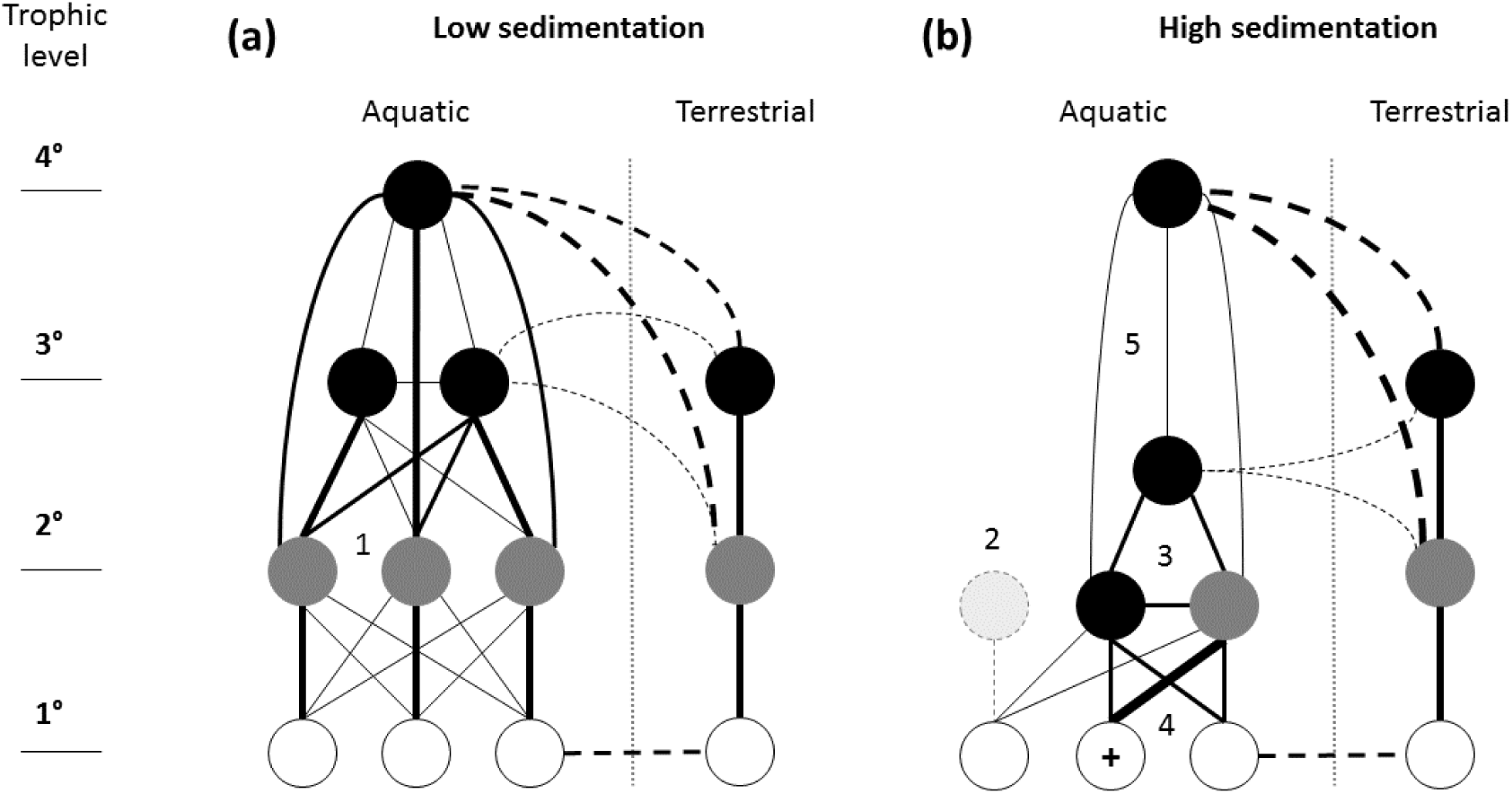
Summary of key food-web changes *potentially* contributing to trophic niche ‘compression’ and other food-web changes associated with sedimentation. Here we have only shown low and high sediment levels to highlight the proposed mechanisms. (a) Low sedimentation allows for (1) niche ‘partitioning’, whereas (b) high sedimentation causes (2) niche ‘elimination’, (3) niche ‘generalists’ with resource scarcity induced omnivory, and under certain circumstances, (4) niche ‘homogeneity’ through ‘hyper-abundant’ resources (+). An additional process involves (5) ‘trophic decoupling’, where top predators maintain trophic position through donor-controlled terrestrial prey subsidies (i.e., unidirectional interactions), but mesopredators track ‘compressed’ aquatic prey, thus leading to a lower trophic position. Basal resources, white; consumers, grey; predators, black. Line weight indicates *possible* interaction strengths; dashed lines, *assumed* trophic pathways from terrestrial food webs.

In contrast to the food scarcity scenario invoked by the niche ‘generalists’ hypothesis, where a resource is ‘hyper-abundant’, it should be particularly attractive to all consumers by virtue of its abundance (Schoener 1974). This is the most likely scenario for the niche ‘homogeneity’ hypothesis (Figs. 1d, 7b). For example, in seagrass ecosystems, trophic diversity decreased as vegetation cover increased, owing to greater isotopic redundancy in consumers and an increased reliance on epiphytic carbon derived from leaves (Calizza et al. 2013). We observed increases in fine benthic organic matter with sedimentation, both in absolute terms and relative to other resources. However, invertebrate ‘collectors’ did not increase numerically to this increased resource base, thus suggesting the niche ‘homogeneity’ mechanism may have been relatively unimportant in our study.

Importantly, the niche ‘elimination’, ‘generalists’ and ‘homogeneity’ mechanisms proposed are not mutually exclusive (Fig. 1), and may interact across trophic levels to impact food-web complexity. This scenario is highly likely in our example, where both the availability of basal resources and communities of prey and consumers were seemingly changing in response to sedimentation. Layman et al. (2007b) found that coastal creek fragmentation led to reduced niche breadth of a resilient generalist top predator, a result they attributed to reductions in the diversity of potential prey (i.e., meaning all predator individuals had to exploit the same resources *sensu* niche ′homogeneity′ as a result of prey niche ′elimination′). Although it was not clear if resource scarcity (i.e., total energy availability) was also a feature of their systems, this example and our study potentially reflect a general ecological principle, where increased dietary specialization and reduced trophic omnivory should occur where habitat conditions allow for an increase in prey and resources that are preferred or optimal (Post et al. 2000). Thus, it seems highly likely that increased omnivory (*sensu* niche ‘generalists’) are important food-web responses to changing prey and resource availabilities caused by disturbances (Wootton 2017).

### Sediment influences on stream fishes

Our community analyses indicated that overall, sediment had weaker effects on stream fishes than benthic invertebrates. Similarly, Yule et al. (2010) observed no influence of sedimentation on fish distributions or diets. The absence of a deterministic influence of sediment on stream fish occupancy patterns highlights how other processes contributing to metacommunity dynamics (e.g., dispersal limitation) may affect the composition of these larger-bodied organisms in freshwaters, because species distributions are typically constrained by network connectivity (Shurin et al. 2009).

Moreover, where fish have generalist-feeding behaviors, terrestrial prey subsidies can offset reductions of benthic prey, thus helping to stabilize stream food webs (Huxel and McCann 1998). For example, Kraus et al. (2015) showed that stream fish increased their use of terrestrial prey due to pollution-induced reductions in sensitive benthic invertebrate prey. In our study, an increased reliance by stream fishes on terrestrial prey subsidies may help explain the lack of a relationship between the whole community δ^15^N range and sedimentation (i.e., by allowing larger fish to retain their trophic position relative to stream invertebrate prey). This may be essential for ‘trophic decoupling’ (Fig. 6,7b), where benthic mesopredators (i.e., *Gobiomorphus* spp.) and their aquatic invertebrate prey became increasingly separated in terms of δ^15^N relative to larger stream fishes (i.e., salmonids and anguilliformes). Future studies should further consider the contribution of resource subsidies to trophic diversity, and when and how these inputs potentially offset *in-situ* prey reductions associated with perturbations to the recipient system.

Lower fish production and reduced abundances are common responses to stream sedimentation (Waters 1995). We were not able to quantitatively measure fish abundances at all sites, but our estimates suggest that declining abundances (*Author′s unpublished data*) may further help explain the non-linear relationship between fish community trophic diversity and deposited sediment. A release from intra- and interspecific competition at high levels of sediment disturbance due to low abundances and diversity may have enabled resident fish to better partition available resources. Larger fish at heavily degraded sites may have foraged over greater distances and relied more strongly on terrestrial resource subsidies, thus acquiring more varied isotopic signatures. This contrasted with the smaller benthic fish seemingly more affected by the proposed ′niche compression` mechanisms related to sedimentation.

### Concluding remarks

We have elucidated food-web responses that highlight the deleterious ecological consequences of landscape disturbance on ecosystems. Although there is evidence that habitat degradation associated with sedimentation strongly contributes to structural changes in invertebrate communities (Burdon et al. 2013), the increasing trophic equivalence of consumers seen here suggests more complex effects on the structural and functional properties of stream food webs. Impaiment to ecosystem functioning might occur because decreasing diversity across trophic levels typically results in lower biomass accrual and resource uptake (Duffy et al. 2007).

However, it is not only species loss, but the functional roles of species that persist which will dictate how food webs and ecosystem functioning responds to environmental change (Layman et al. 2007b, Lindo et al. 2012). One consequence may be decreased stability, because trophic niche ‘compression’ indicates that energy flow paths are more homogenous, potentially leading to less stable food-web structures (Rooney et al. 2006, Layman et al. 2007b). This may be particularly profound where landscape disturbances alter the stabilising influence of trophic interactions with differing strengths and distributions (Wootton and Stouffer 2016). While it seems likely that terrestrial resource subsidies help offset changes for higher trophic levels in our streams, trophic simplification may still render top predators more susceptible to population fluctuations and local extinctions (Layman et al. 2007b). With ecosystems facing multiple anthropogenic threats globally, understanding the consequences of environmental change requires ecologists to apply novel methods that provide greater insights into the functional consequences of biodiversity loss. This knowledge will ultimately contribute to the improved conservation of ecosystems and successful remediation of degraded habitats.

## Acknowledgements

We thank landowners for allowing property access. Richard White and Duncan Gray helped with fieldwork, Hayley Devlin contributed in the laboratory, and Linda Morris provided technical support. Members of FERG gave useful feedback. Andrew Jackson and attendees of the PR∼Statistics Workshop ‘Analysis of Stable Isotope data using SIAR’ at Rowardennan, Scotland (27-30^th^ July 2015) provided stimulating discussions. F.J.B. was a scholarship recipient from the Mackenzie Charitable Foundation, and the Brian Mason Scientific and Technical Trust provided additional financial support for isotope analyses. Fish and crayfish were collected under the University of Canterbury Animal Ethics Permit 2009/37R.

## Author′s contributions

F.J.B. and J.S.H. conceived the research; F.J.B. created the conceptual framework, designed the research, collected the data, conducted analyses, and wrote the paper. All authors contributed critically to the draft and gave final approval for publication.

## Data accessibility

Data will be made available via the Dryad Digital Repository.

## Supporting Information

**Appendix A**

Study sites

**Appendix B**

Detailed methods descriptions

**Appendix C**

Additional results

